# Compartmentation of Putrescine Synthesis in Plants

**DOI:** 10.1101/2022.09.03.506421

**Authors:** Kumud Joshi, Sheaza Ahmed, Lingxiao Ge, Vipaporn Phuntumart, Andrea Kalinoski, Paul F. Morris

## Abstract

Three plant pathways for the synthesis of putrescine have been described to date. These are the synthesis of putrescine from ornithine, by ornithine decarboxylase (ODC); and the synthesis of putrescine from arginine by arginine decarboxylase, agmatine iminohydrolase (AIH) and N-carbamoylputrescine amidohydrolase (NLP1); or arginine decarboxylase and agmatinase. Several enzymes associated with putrescine synthesis have yet to be localized. Here we showed that ODC in soybeans and rice was localized to the ER. In rice, agmatinase is localized to the mitochondria. In *A. thaliana* there are five isoforms of AIH and three isoforms of NLP1. Stable GFP-tagged transformants of the longest isoforms of AIH and NLP1 showed that both proteins were localized to the ER in leaves and roots of *A. thaliana*. Four of the isoforms of AIH and all of the isoforms of NLP1 were localized to the ER. However, AIH1.4 was localized to both the ER and the chloroplast. Combining these results with other published data, reveal that putrescine synthesis is excluded from the cytoplasm and is spatially localized to the chloroplast, ER and likely the mitochondria. Synthesis of putrescine in the ER may facilitate cell to cell transport via plasmodesmata, or secretion via vesicles. Differential expression of these pathways may enable putrescine-mediated activation of hormone-responsive genes.

## Introduction

Polyamines are ubiquitous polycationic molecules found across all domains of life. In plants, polyamines play a role in growth, development, and stress responses. Polyamines confer tolerance against multiple ranges of abiotic stresses such as drought stress, heat stress, salt stress, cold/freezing stress, flooding stress, and biotic stresses in plants (Alcázar& Tiburcio 2014; Juan F. Jiménez-Bremont et al.,2013). Changes in plant polyamine levels influence different functions such as transcription, RNA modification, protein synthesis, enzyme activity, and signaling pathways involving jasmonic acid, salicylic acid, abscisic acid, gibberellin, and Nitric Oxide (Kusano et al.2008; Kusano et al. 2007; Evans et al.,1989; Galston et al.,1997; Takahasi et al., 2010; Wimalasekera et al., 2011; Sagor et al.,2015). They are also involved in different ranges of other biological processes such as senescence, protection from different types of environmental stresses, protection from infections such as viruses and fungi, etc. (Tiburcio et al.,1993; Galston et al.,1997; Bais and Ravishankar 2002).

The synthesis of putrescine, the smallest of the polyamines can occur via several routes. The direct conversion of ornithine to putrescine is achieved by ornithine decarboxylase (ODC) however, ODC is absent in Brassicaceae (Hanfrey et al., 2001) the moss genome (Hanfrey et al., 2001) and in the duckweed *Spirodela polyrhiza* (Upadhyay et al., 2021). The second pathway for putrescine synthesis involves the synthesis of putrescine from arginine by a three-step process initiated by arginine decarboxylase-1 (ADC1) and the sequential activity of agmatine iminohydrolase (AIH), and N-carbamoyl putrescine amidohydrolase (NLP1) (Janowitz et al., 2003).

The third pathway results from the conversion of agmatine to putrescine by arginase/ agmatinase (ARGAH) enzymes (Patel et al 2017). In *A. thaliana*, ADC2 and arginase/agmatinase ARGAH2 are localized to the chloroplast (Patel et al., 2017). All plant arginase/agmatinase enzymes are very structurally conserved and have both arginase and agmatinase activity (Patel et al.,2017; Sekula 2020). A unique feature of plant ARGAHs is that they form hexameric complexes, and substrate binding of ornithine or agmatine is stabilized by a loop from the neighboring subunit (Sekula 2020). ARGAH enzymes have historically been categorized only as enzymes that catalyze the catabolism of arginine to ornithine. Seed proteins are rich in arginine and during germination, proteolysis of seed proteins and catabolism of arginine by ARGAH provides nitrogen and carbon for the synthesis of other amino acids (Siddapa and Marathe 2020). However, in plant leaves, arginine is only a minor amino acid (Tan et al., 2010; Shi et al 2013) and the K_m_ of ARGAH for arginine is 30 mM (Chen et al., 2004), while the K_m_ for agmatine is 73±10 μM (Patel et al., 2017). When ADC2 and ARGAH2 were mixed with 1mM arginine, only agmatine and putrescine were found in significant amounts (Patel et al., 2017). Moreover, when ADC2 and ARGAH enzymes were co-expressed in conditionally lethal *spe2* yeast cells (deficient in putrescine synthesis) these enzymes supported strong and rapid growth of yeast relative to WT cells. This would not have been observed had ARGAH been competing with ADC2 for arginine. Total arginine levels were also not significantly affected by either AtARGAH knockout or overexpressing lines (Shi et al. 2013) which might be expected given the low substrate affinity of the enzyme for arginine. Many plant genomes have two ADCs (Patel et al., 2017). The differential activation of plant ADCs may enable the production of agmatine to be metabolized by different enzymes (AIH and NLP1, or ARGAH) that are in different plant compartments.

In *A. thaliana*, two enzymes enable the directed synthesis of acetyl-putrescine. N-acetyl transferase 1 (NATA1) converts ornithine to N^δ^-acetyl ornithine while ADC1 converts N^δ^-acetyl ornithine to acetyl-putrescine (Lou et al. 2020). NATA1 also synthesizes acetyl-putrescine, but its affinity for ornithine is higher.

Plants have an absolute requirement for putrescine synthesis (Tabor &Tabor 1964). The observation that homozygous AIH mutants have a defective embryo phenotype was originally interpreted as evidence that *A. thaliana* and other Brassicas had only a single putrescine synthesizing pathway (Tzafrir et al., 2004). However, homozygous mutants of *nlp1*do not exhibit a deleterious phenotype under normal growing conditions, thus indicating that putrescine can be synthesized by more than one pathway in the model plant. Polyamines produced from different pathways have different roles in various plant functions (Upadhaya et al 2021). In tomato leaves, SlODC*2* was significantly upregulated in response to drought and salt, while other putrescine synthesizing enzymes were unchanged or downregulated. Polyamines produced via the ODC pathway promote flowering whereas polyamines derived from the ADC pathway are involved in vegetative development (Kakkar &Swahney 2002). Additionally, ODC plays role in cell division and proliferation whereas ADC levels are up-regulated in stressed tissue and during cell extension (Perez-Amador et al., 1995; Soyka and Heyer 1999). In *Solanum lycopersicum* and *Datura stramonium*, ODC expression is very low in mature leaves whereas it is upregulated in the roots and stem (Perez-Amador and Granell.,1995). In soybeans, both ADC and ODC genes are upregulated in the hypocotyl and root tips indicating a co-expression pattern during some physiological events such as organ development and cell expansion (Delis et al., 2005).

Plants may have one or two agmatinase. GFP-tagging experiment have localized one soybean and one *A. thaliana* agmatinase to the chloroplast (Patel et al.2017). But proteomic data also indicates that ARGAH is present in the mitochondria of *A. thaliana* (Senkler et al., 2017). Thus, this data suggests that putrescine synthesis can take place in chloroplast and mitochondria if agmatine can be imported into mitochondria.

Crosstalk regulation of putrescine synthesis may exist between different putrescine pathways. In homozygous mutants of *Arabidopsis*, *argah1* and *argah2* strains show an increased level in the expressions of the *AIH* and *NLP1* genes along with *ADC1* and *ADC2* genes, thus an increase in putrescine level (Shi et al., 2013). Contrary to this, *AtARGAH1* and *AtARGAH2* overexpressing lines showed a decrease in the levels of all four genes as compared to the wild type and a decrease in putrescine levels suggesting an upregulation of the *ADC1* pathway (Shi et al., 2013).

In plants, alternative splicing occurs in up to 70% of intron-containing genes (Chaudhary et al., 2019). Thus, alternative splicing of pre-mRNA is an important step for gene regulation (Shang et al.,2017) The number of variants differ depending on the tissue type, environmental conditions, and developmental stages (Syed et al., 2012). Different environmental conditions can alter the expression of alternatively spliced gene isoforms. For example, in potatoes, the cold-induced sweetening phenotype results from the conversion of starch to glucose and fructose by invertase. Potato lines that are resistant to cold-induced sweetening exhibit higher expression levels of two invertase inhibitor isoforms (INH2a and INH2b) relative to that seen in susceptible lines (Brummell et al., 2011).

Alternative splicing can result in targeting of proteins to different subcellular compartments. Knowing the subcellular location of a protein in a cell is an important component of understanding its functions (Andrade et al., 1998). Auxin biosynthesis occurs *via* multiple pathways in plants. (Kriechbaumer et al., 2011) show that the two isoforms of YUCCA4, an auxin biosynthetic gene are tissue-specific and are differentially localized to the ER and the cytoplasm. One isoform is present only in flowers and is anchored to the cytosolic face of the endoplasmic reticulum membrane. The other isoform is found in all tissue types and at a cellular level distributed throughout the cell. Alternative splicing of Transthyretin-like (TTL) protein results in the loss of an internal peroxisomal targeting signal and shifts the resulting protein product from the peroxisome to the cytosol (Lamberto et al., 2010).

Compartmentation is an essential feature of plant metabolism. Here we have addressed where key enzymes involved in putrescine synthesis are located using confocal imaging of transiently expressed genes in *N. benthamiana* and stable transformations in *A. thaliana*. We note that putrescine synthesis is specifically excluded from the cytosol, and the major sites for putrescine synthesis are the chloroplasts, ER and mitochondria.

## Materials and Methods

### Plant Growth Conditions

#### Arabidopsis thaliana

*A. thaliana* Col-0 ecotype was used as the wild type. *AIH* and *NLP1* over-expression lines were generated as per the protocol by (Tang 2012; mtu.edu/~gtang1/Tang_Website/Lab_Protocols). Briefly, *A. thaliana* seeds were surface sterilized by mixing about 120 seeds in a 1ml mixture of 0.1% Triton X and 10% bleach. The seeds were then rinsed with sterile water 3 times before placing them in the refrigerator at 4 degrees for stratification. After stratification, the plants were transferred to soil in 4-inch pots. Plants were grown under (16 h light intensity 110 μmol m−2 s−1 and 8 h dark) at 22°C ± 1°C.

To prevent the seeds from drying out before the germination was established, plants were covered with transparent plastic wrap with holes for 5-7 days.

#### Nicotiana benthamiana

*N. benthamiana* seeds were planted in a germination mix (Lambert LM-GPM) in a tray by carefully placing seeds ½ inches apart and covering them with a small layer of the potting mix. After germination (10-14 days), seedlings were transplanted into packs or pots of Lambert AP all-purpose) potting mix and grown in the greenhouse where temperatures were maintained between 18-30° C. Natural light in the greenhouse was supplemented during short daylength periods and cloudy days for an average photoperiod of 12-14 hours. Plants were watered as needed by soaking the bottom of the pot.

### DNA source and constructs

cDNA clones of *AtAIH.1* (At5G01870) and *AtNLP1.2*(At2G 27450.2) were obtained from Arabidopsis Biological Research Center, Ohio State University, Columbus, Ohio. Sequences of *GmODC OsODC* and *OsAg*, *AtAIH-4, AtNLP1-1*, and *AtNLP1-3* were codon-optimized and synthesized by Genscript, NJ, and Thermo Fisher, MA. The mCherry-ER marker (ER-rbCD3960) used in this study was obtained from Arabidopsis Biological Resource Center. Full-length sequences of *GmODC*, *OsODC*, *OsAg*, *AtAIH.1*, *AtAIH.2*, *AtAIH.4*, *NLP1.2*, *NLP1.1*, and *NLP1.3* were cloned into plant expression vectors pGWB6, or pGWB5 using the GATEWAY® recombination system (Nakagawa et al., 2007). Genes were amplified using gene-specific primers listed in **Supplemental Table 1** following the manufacturer’s protocol for **PrimeSTAR** GXL DNA polymerase. Agarose gel electrophoresis was performed to confirm the PCR products. Topo cloning reaction was performed using the pENTR™/SD/D-TOPO® Cloning Kit (Thermo Fisher, MA) as follows: Fresh PCR product 0.5–4 μL; 0.5–4 μL Salt solution;1 μL — Dilute salt solution (1:4) — 1 μL Water to a final volume of 5 μL. The solution was mixed gently and incubated for 5 mins at room temperature. The reaction was then placed on ice and transformed into chemically competent Top10 cells by heat shock transformation method. 2 μL of the TOPO® Cloning reaction was added to a tube of One Shot® chemically competent *E. coli* cells and mixed gently. The tube was then incubated on ice for 5–30 minutes. Cells were then heat shocked for 30 seconds at 42°C by placing them in a water bath without shaking. The tube was then immediately placed on ice. S.O.C. medium)250 μL was added to the tube and incubated at 37°C for 1 hour with shaking. The aliquots of 50–200 μL of bacterial culture were then spread on a prewarmed selective LB plate (Kanamycin) and incubated overnight at 37°C. Positive colonies were selected and cultured LR reaction was performed using LR Clonase™ II enzyme mix (Thermo Fisher, MA) following the manufacturer’s protocol with a modification of the incubation time (3h instead of 1h). The cells were plated onto agar plates with appropriate antibiotic selection (Kanamycin/Hygromycin). Positive colonies were selected and cultured. Plasmid extraction was done using the Zippy Plasmid Miniprep Kit (Zymo Research, CA). Plasmid DNA from this step was used in the next step for *Agrobacterium* transformation.

### Transient expression in *Nicotiana benthamiana*

Chemically competent Agrobacterium cells of *Agrobacterium tumefaciens* strain GV3101 were prepared using the freeze- thaw method (Xu and Li 2008). Briefly, one μg of plasmid DNA (respective constructs) was added to 0.1ml partially thawed chemically competent *Agrobacterium* cells and mixed gently. The tube was then placed in liquid nitrogen and allowed to be frozen. The frozen cells were then thawed in a 37°C water bath for 5 minutes. SOC medium (150 uL) was then added to the thawed cells and incubated at 28°C for 2-4hrs with gentle shaking. The cells were plated on LB plates containing Kanamycin (50ug/ml)/Hygromycin (50 μg/ml)/Gentamycin (10 μg/ml) and incubated at 28°C for 36 hours. The positive colonies were used to culture and infiltrate tobacco leaves.

*Agrobacterium* transformed with genes of interest were cultured in 2ml LB with appropriate antibiotics at 28°C 150 rpm for 24-36hrs. The overnight culture (1ml) was then transferred to an Eppendorf tube and centrifuged for 5 minutes at 5000RPM. The supernatant was discarded, and the pellet was resuspended in 1ml infiltration solution (10mM MgCl_2_ and 100μM acetosyringone). Cells were washed and resuspended twice in infiltration solution. The final pellet was then suspended in 200μL of infiltration solution. The cell suspension was adjusted to OD600 of 0.5 for co-infiltration studies. Finally, 1ml of the cell suspension containing gene construct and 1ml of the cell suspension containing marker was mixed and infiltrated to the lower surface of the 5-6 weeks old leaves of tobacco on either side of the major vein (Sparkes et al., 2006).

### Stable transformation via floral inoculation method

AIH and NLP1 transgenic plants were generated via “the floral inoculation method” as per the protocol by (Tang 2012). Briefly, Arabidopsis seeds were sterilized and stratified for 3 days at 4°C. *Agrobacterium* strains containing the gene of interests were inoculated into a 5ml liquid LB medium containing appropriate antibiotics (Kanamycin, hygromycin, gentamycin). The cultures were then incubated at 28°C for 2 d. The feeder cultures were used to inoculate a 500ml liquid LB with appropriate antibiotics. The cultures were then grown at 28°C for 16–24 h until the cells reached an OD600 no less than 1.

*Agrobacterium* was harvested by centrifugation for 20mins at room temperature at 5500rpm; supernatant poured, and remaining media removed with a pipet as much as possible. Then, the cells were then re-suspended in the inoculation medium (5% sucrose). and mixed along with 5μl of Silwet L-77(Tang 2012).

For the modified floral inoculation procedure, 50μL of *Agrobacterium* cells in 5% sucrose solution was added to every flower bud. The flowerpots were then placed in a black box for three days and then returned to the growth chamber to normal growth conditions. The inoculation procedure was repeated a week later on newly emerging buds. These transgenic T1 seeds were collected after 3 weeks, then planted using the same method as mentioned above. After 5 days, seedlings were sprayed with Basta every 2-3 days for selection. Healthy green plants were then transplanted and leaves from 2–3-week-old plants were harvested for analysis.

### Confocal Analysis

Images were acquired from different infiltrated leaf sections of tobacco at 12-72 h after leaf inoculation using a Leica TCS SP5 laser scanning confocal microscope (Leica Microsystems, Bannockburn, IL). Images were acquired in the XYZ plane in 0.25 μm, 0.5 μm or 1 μm steps with a 63X oil objective (NA 1.40) using the sequential scan mode to eliminate any spectral overlap in the individual fluorophores. GFP was excited at 488 nm and detected at 510 nm. mCherry was excited at 561 nm and detected at 610 nm. Plastids were excited at 633 nm and detected at 670 nm. GFP signals were false-colored green, mCherry fluorescence was false-colored red, and chlorophyll autofluorescence was false-colored blue. Background fluorescence from untransformed leaves of plants was determined using the same acquisition settings and subtracted from the images to identify fluorescence generated by tagged proteins. ImageJ was used to merge the images (Hartig et al., 2013). Images and videos of image Z-stacks, taken as planar sections from transformed *N. benthamiana* leaves, were made using the Leica Application Suite Advanced Fluorescence (LASAF) program.

### Upstream Open Reading Frame analysis

To analyze 5’ regulatory upstream untranslated regions preceding the open reading frames (uORF), of the AIH and NLP1 genes, the sequences were obtained from TAIR. Presence of uORFS in the 5’ region of the genes was determined using National Library of Medicine- ORF finder tool (https://www.ncbi.nlm.nih.gov/orffinder/).

## Results

### Ornithine decarboxylase is localized to the ER

The localization of ODC in plants has not yet been established. Plant genomes may have more than one ornithine decarboxylase (Sivakumar et al., 2021). The soybean genome contains two predicted gene models for ornithine decarboxylase (Glyma04g020200.1 and Glyma06g02290). These soybean sequences share 93% identity of amino acid residues, with strong conservation of sequence identity at both N- and C-terminal regions (Phytozyme). To identify the intracellular location of Glyma_04g020200.1 (GmODC) we generated C- terminal GFP-fused protein constructs and performed an analysis by confocal microscopy. The GFP fusion of Glyma_04g020200.1 and a mCherry ER marker was used to identify cells expressing both markers (Figure 1, A& B). Chlorophyll fluorescence was used as a marker to localize the chloroplast within mesophyll cells and was false colored blue (**Figure 1, C**). Transformed cells showed that GmODC was localized to discrete regions of the cytoplasm that overlapped with the mCherry ER marker **(Figure 1, D)**. In transformed mesophyll cells, analysis of individual frames of the stacked images showed that the GmODC was localized within the cortical networks and sheets, of the ER as the GFP signals overlapped with those of mCherry ER marker **(Movie.S1)**. The subcellular localization of Os02G0482400 (OsODC) was also determined by transient expression of N- and C-terminal GFP fusions in *N. benthamiana* leaves mediated by *Agrobacterium* (**Figure 2**). The GFP signals overlapped with the mCherry signals of the endoplasmic reticulum (ER) marker. These results suggest that OsODC is also localized to the ER. Both N-terminal and C-terminal GFP fused constructs were generated and co-localized with ER markers (**Movie. S2)**. We noted that N-terminal fused GFP localization (**Figure 2: (a-d** was more prominent than C- terminal constructs (**Figure 2: e- h)**. Thus, at least one of the soybean genes is localized to the ER and in rice where there is only one gene and no alternatively spliced form, the ODC protein is localized to the ER

**Figure 1.**
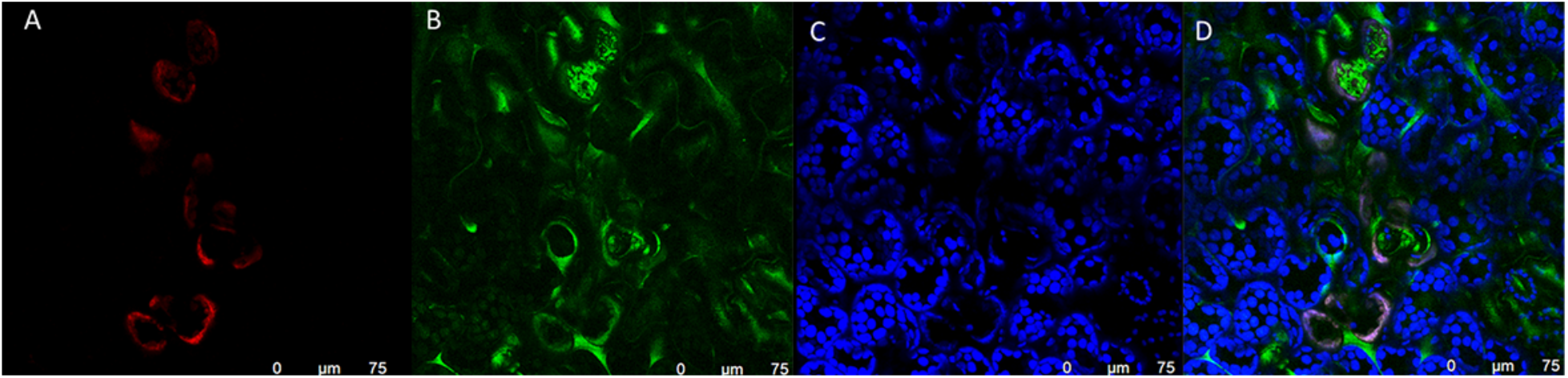
Subcellular localization of GmODC-GFP in the Endoplasmic Reticulum of mesophyll cells of *N. benthamiana*. GmODC-GFP and ER mCherry were expressed in *N. bethamiana* leaves and shown here as Z-stacked confocal images. (A) ER-mCherry marker (B) GFP GmODC (C) Chlorophyll autofluorescence has false color blue (D) merged image. In cell layers where GFP-GmODC overlap with mCherry, the images are pink. Images were taken 3 days after the infiltration of *Agrobacterium* into tobacco leaves and were background subtracted. Scale bars, 75μm.

**Figure 2.**
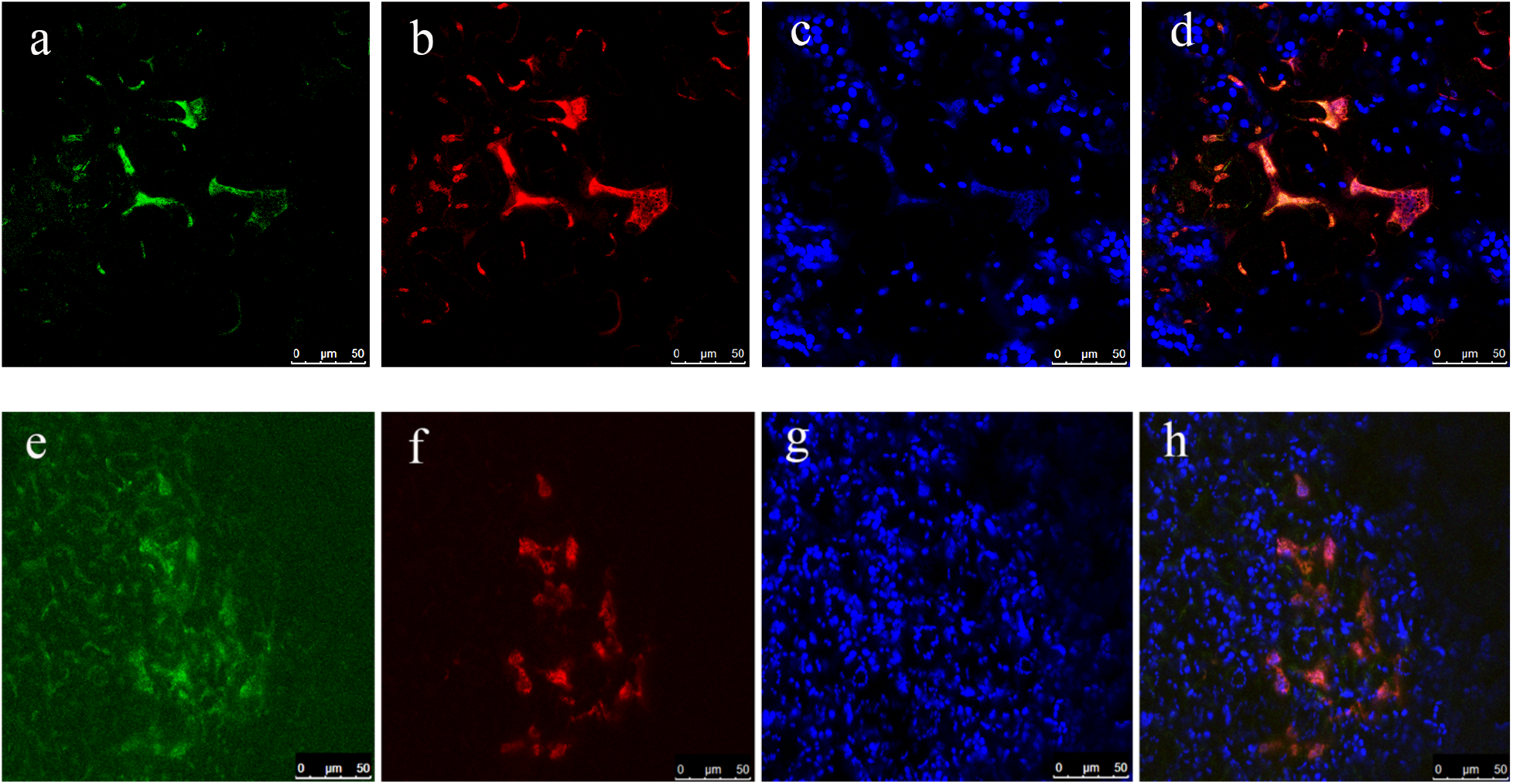
Subcellular localization of GFP- OsODC in the Endoplasmic Reticulum of mesophyll cells of *N. benthamiana*. N-terminal GFP-OsODC fusions are shown in and d. C-terminal OsODC-GFP are shown in e, h. GFP-OsODC and ER mCherry were expressed in *N. bethamiana* leaves. a, GFP-OsODC; e, OsODC-GFP; (b, f) ER-mCherry marker (c, g) Chlorophyll autofluorescence has false color blue (d, h) merged image. In cell layers where GFP-OsODC or OsODC-GFP overlap with mCherry, the images are pink. Images were taken 3 days after the infiltration of Agrobacterium into tobacco leaves and were background subtracted. Scale bars, 50μm.

### Rice arginase/agmatinase (OsARGAH) is localized to the mitochondria of leaves

Mitochondrial localization of OsARGAH in the root hypocotyl of transgenic rice has already been demonstrated by (Ma et al., 2013). The N-terminal of this gene contains a predicted mitochondrial signal peptide (data analyzed by PSORT, Signal P (Petersen et al., 2011). To address the localization of this protein in photosynthetic tissue, we expressed OsARGAH-GFP in *N. benthamiana* and determined its localization by confocal microscopy. Examination of the Z-stack showed a punctate distribution of GFP fluorescence throughout the cell with some areas overlapping the chloroplast (**Figure 3, a, b, c**). However, examination of representative optical planes within the Z-Stack (showed that organelles labelled green were not associated with the chloroplasts (**Figure 3, f**). The punctate localization pattern in these images is similar to previous report for other GFP-tagged mitochondrial proteins (Wolter et al. 1997). This localization pattern is also consistent with the proteomic data of purified mitochondria from rosette leaves of *A. thaliana* (Senkler et al. 2017). We conclude that OsARGAH is localized to mitochondria. Thus, the mitochondria can also be a site for putrescine synthesis, if agmatine is imported and converted to putrescine by OsARGAH.

**Figure 3.**
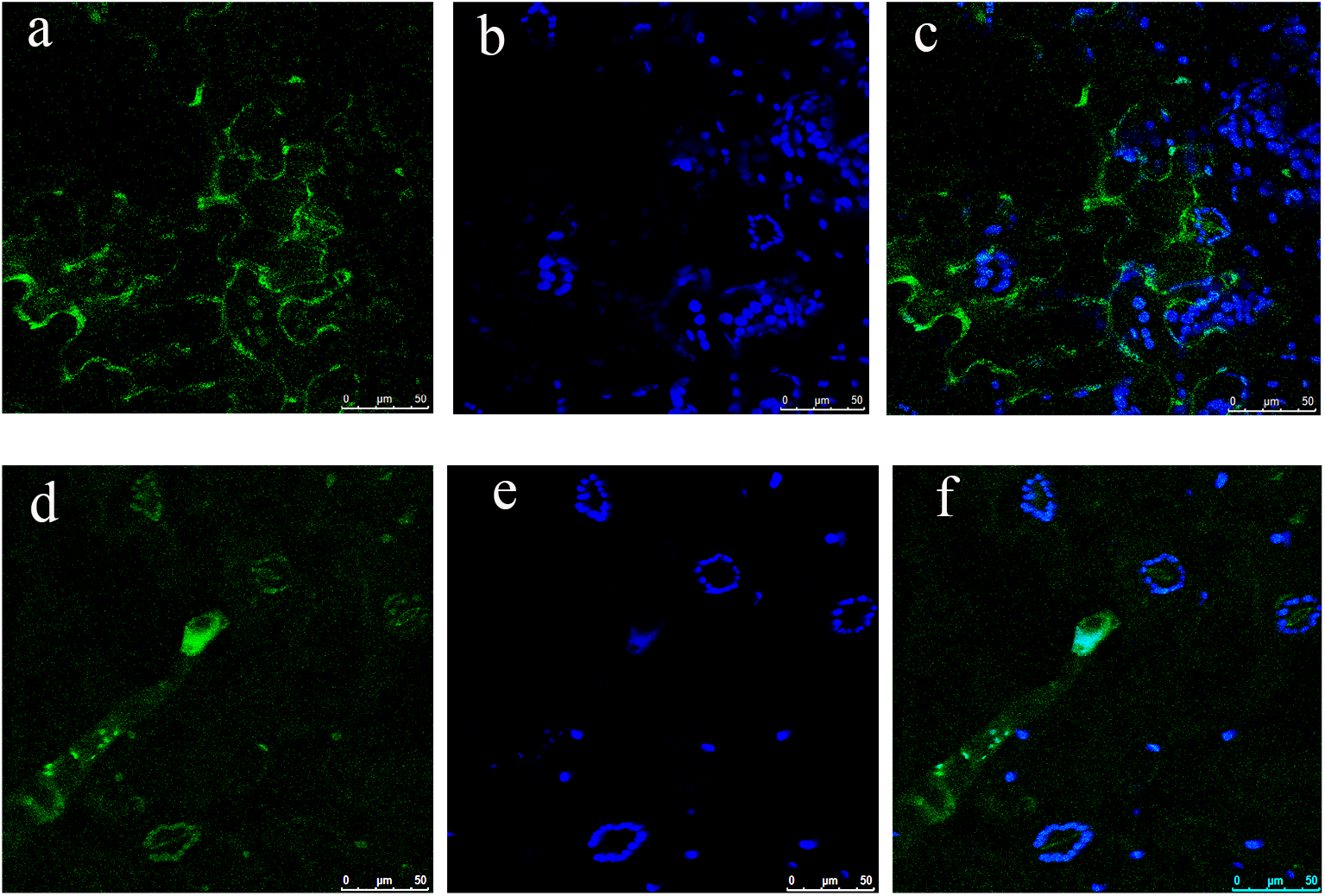
OsARGAH-GFP is localized to the mitochondria in mesophyll cells of *N. benthamiana*. GFP- fused constructs were infiltrated in the plant cells by *Agrobacterium* infiltration. GFP signals were localized to the mitochondria. **Figure (a, b, c-Z-stack) and Figure (d, e, f- Z- slice** (a, d) Transient expression of OsARGAH-GFP, green (b, e) Chlorophyll autofluorescence false coloured blue (c, f) Merged image. Images were taken 3 days after the infiltration of Agrobacterium into tobacco leaves and were background subtracted. Scale bars, 50μm.

### Agmatine iminohydrolase and its isoforms are localized in the Endoplasmic reticulum and the chloroplast

In plants, one of the pathways for putrescine synthesis occurs by the conversion of arginine via agmatine and N-carbamoyl putrescine to putrescine via the action of ADC1, AIH, and NLP1. This pathway was described twenty years ago by (Janowitz et al., 2003) in *A. thaliana*. In silico analysis showed that there are five splice variants of *AIH* in *A. thaliana* (**Figure S1.1**.). The full-length of putative AIH.1 protein is 383AA long and the splice variants AIH.2; AIH.3 AIH.4 AIH.5 produce proteins of 299, 299, 377, 299 amino acids, respectively (Berardini et al., 2015). All five splice forms share the same seven, C-terminal exons but differ in the transcriptional start sites, 5’ UTRs, and the inclusion of different N-terminal exons. The splice variants, 2,3, and 5 differ in the length of the 5’ UTRs but share the same coding sequence. Variants 1 and 4 share two additional exons in common in the N-terminal but they have different N-terminal start sites and processing thus resulting in a unique N-terminal peptide sequence. The presence of multiple AIH isoforms suggests that they are differentially regulated.

To identify the subcellular localization of AIH in plants, we generated C-terminal GFP constructs for each of the three distinct putative proteins of AIH (383 aa, 299aa, 377aa) and transiently expressed them in the leaves of *N. benthamiana*. We examined both individual 0.5μm optical slices, Z- stacked images, and movies of single and merged images. Two samples of AtAIH.1-GFP and mCherry ER are shown in (**Figure 4**). In both cases, stacked images show AIH to be discretely localized within the cytoplasm, and this fluorescent pattern co-localized with the mCherry ER marker (**Movie S3)**. Next, we examined the localization of AIH.2-GFP in a Z-stack and an optical section within Z-stack. In each case, the fluorescence of AIH.2 was intensely localized to the ER cisternae (**Movie S4)**. Merged image of this Z stack with a bright field is shown in (**Figure 5**).

**Figure 4.**
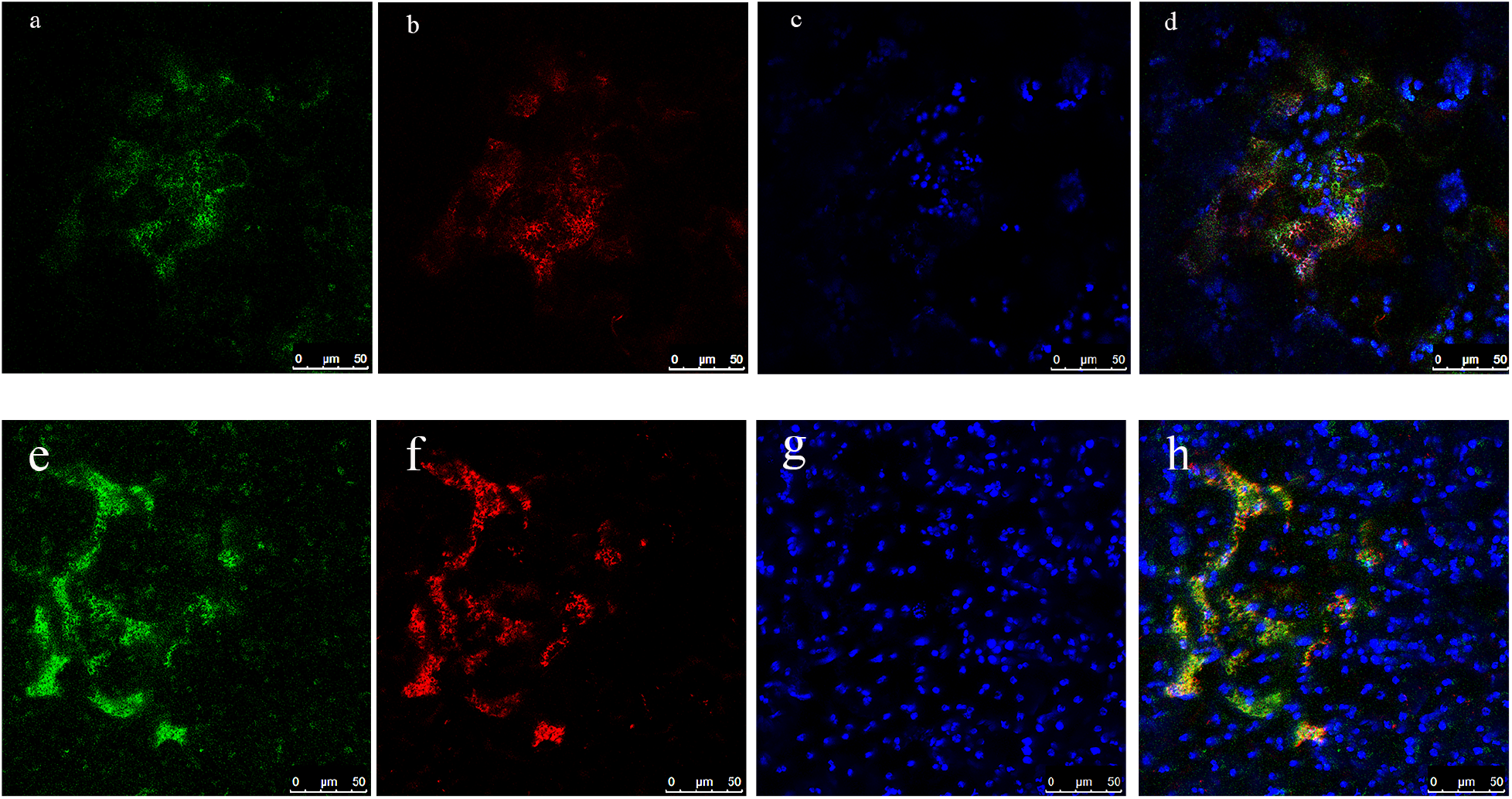
Localization of AtAIH.1-GFP in the mesophyll cells of *N. benthamiana*. Z-stacks of two regions from different leaves are shown. (a, e) AtAIH-GFP (b, f) m-Cherry ER marker (c, g) Chlorophyll auto fluorescence false-colored blue (d, h) merged images. Scale bars 50μm. 3D visuals of the localization presented as a movie in the supplemental section.

**Figure 5.**
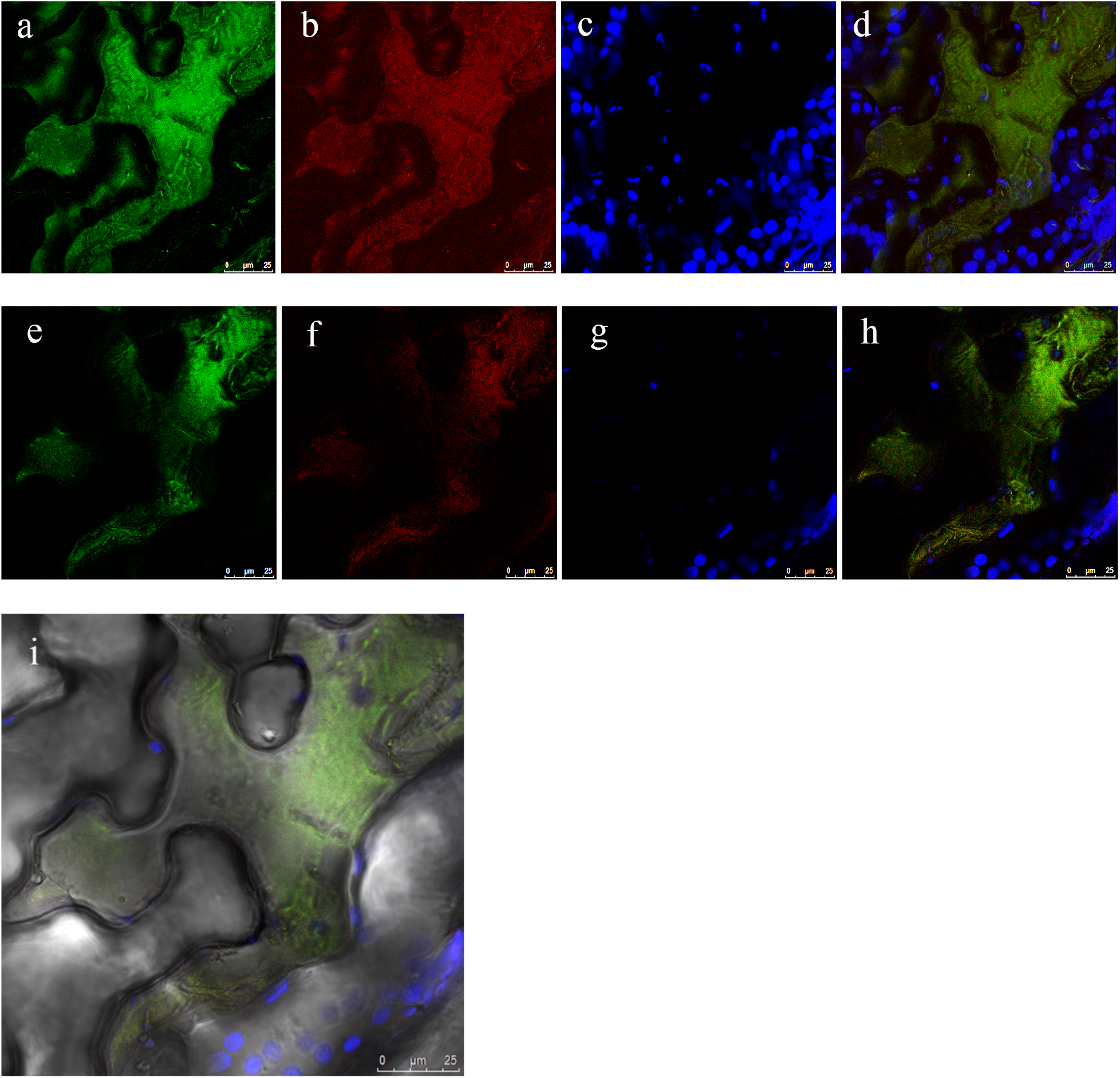
Subcellular localization of AtAIH.2 in the mesophyll cells of *N. benthamiana* with GFP fused protein along with an ER mCherry organellar marker showing ER localization of the gene. Figure (a, b, c, d- Stacked images) & Figure (e, f, g, h, slice) (a, e) AtAIH-GFP (b, f) m-cherry ER marker (c, g) Chlorophyll auto fluorescence, false-colored blue (d, h) Merged Image. Figure (i): Brightfield Image showing GFP signal in the ER. Scale bars 25μm.

We next examined the localization of the isoform AIH.4 in *N. benthamiana* (**Figure 6**). Examination of GFP fluorescence patterns in transformed cells (**Figure 6)**. In merged images, most of the expression of was associated with the chloroplast (**Figure 6 d)** but in the optical slices, some of the fluorescence also appeared to be localized to the ER (**Movie S5)**. Notably, the protein localization tool Plant-mPLOC also predicts that this protein is localized to the chloroplast (Kuo-chen et al 2010).

**Figure 6.**
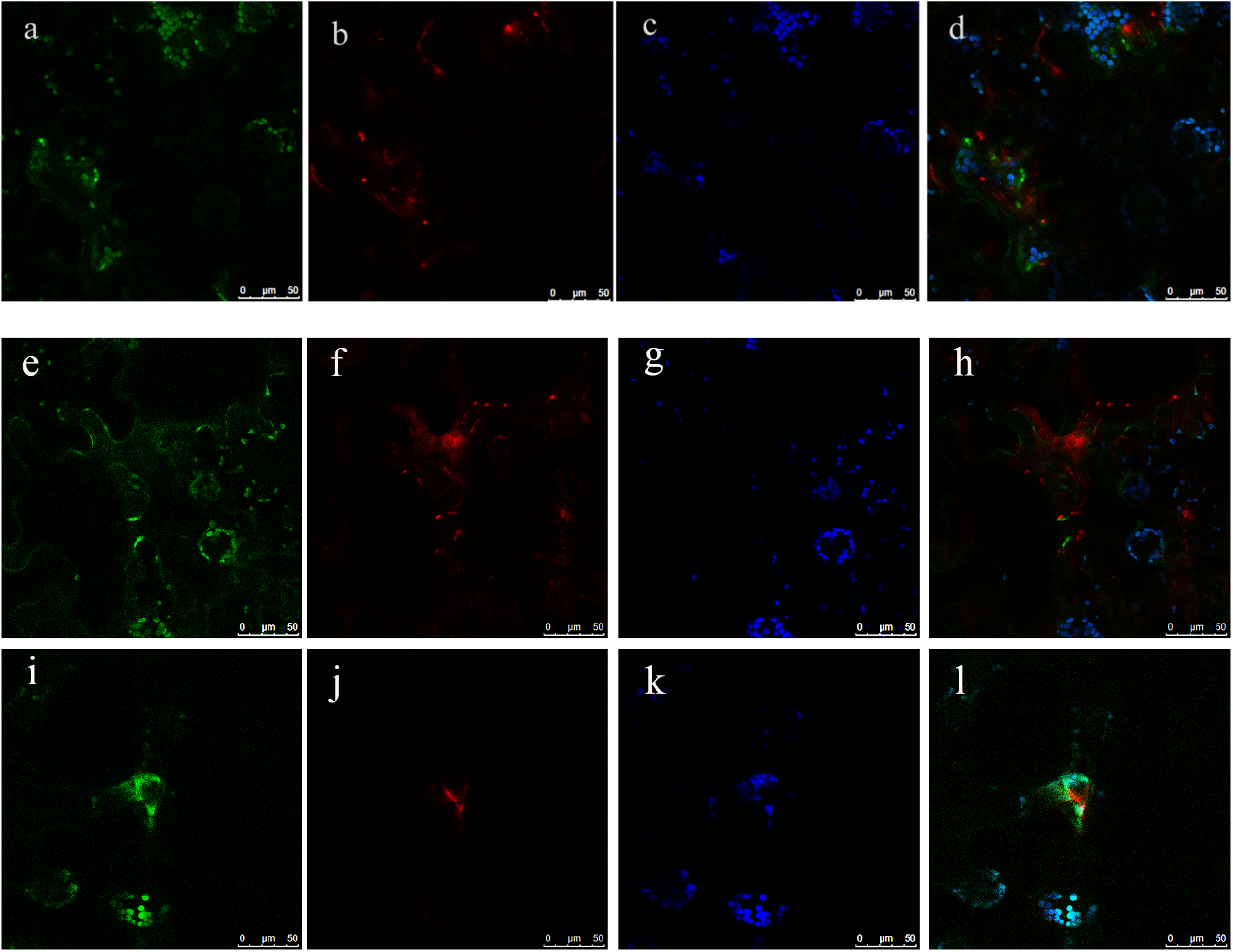
Subcellular localization of AtAIH.4-GFP in the mesophyll cells of *N. benthamiana.* AIH.4 was co-transformed with an ER mCherry organellar marker. Figure (a, b, c, d- Stacked images) & Figure (e, f, g, h, I, j, k, l- single optical slice) highlighting chloroplast and ER GFP signals in different areas of the leaf tissue (a, e, i) AtAIH-GFP (b, f, j) m-cherry ER marker (c, g, k) Chlorophyll auto fluorescence (d, h, l) Merged image indicates co-expression of AIH.4 in the ER and the chloroplast. Scale bars 50μm.

To further address the localization of AIH.1, we generated overexpression lines of AIH.1 in Arabidopsis and used confocal microscopy to examine both leaves and roots (**Figure 7 & Figure 8**). Examination of optical slices of the leaf images shows that the GFP signal only partially overlaps with that of thylakoids. Other areas were strongly associated with GFP fluorescence distinct from the chloroplast. Taken together, this fluorescent pattern is consistent with localization in the ER membranes abutting the chloroplast **(Figure 7; Movie S6**). The contact between ER-chloroplast membranes is not uncommon in plants and is essential for plant homeostasis and lipid biosynthesis (Liu and Li., 2019). The ER- Chloroplast contact sites are involved in the trafficking of proteins and lipids. Lipid transporters such as trigalactosyldiacylglycerol (TGD) exclusively localize to the ER-Chloroplast contact sites in *Brassica napus L* (Tan et al., 2011). In roots, the ER signals form a network of fluorescence throughout the cells consistent with the spatial expanse of the ER in plant cells (**Figure 8; Movie S7**).

**Figure 7.**
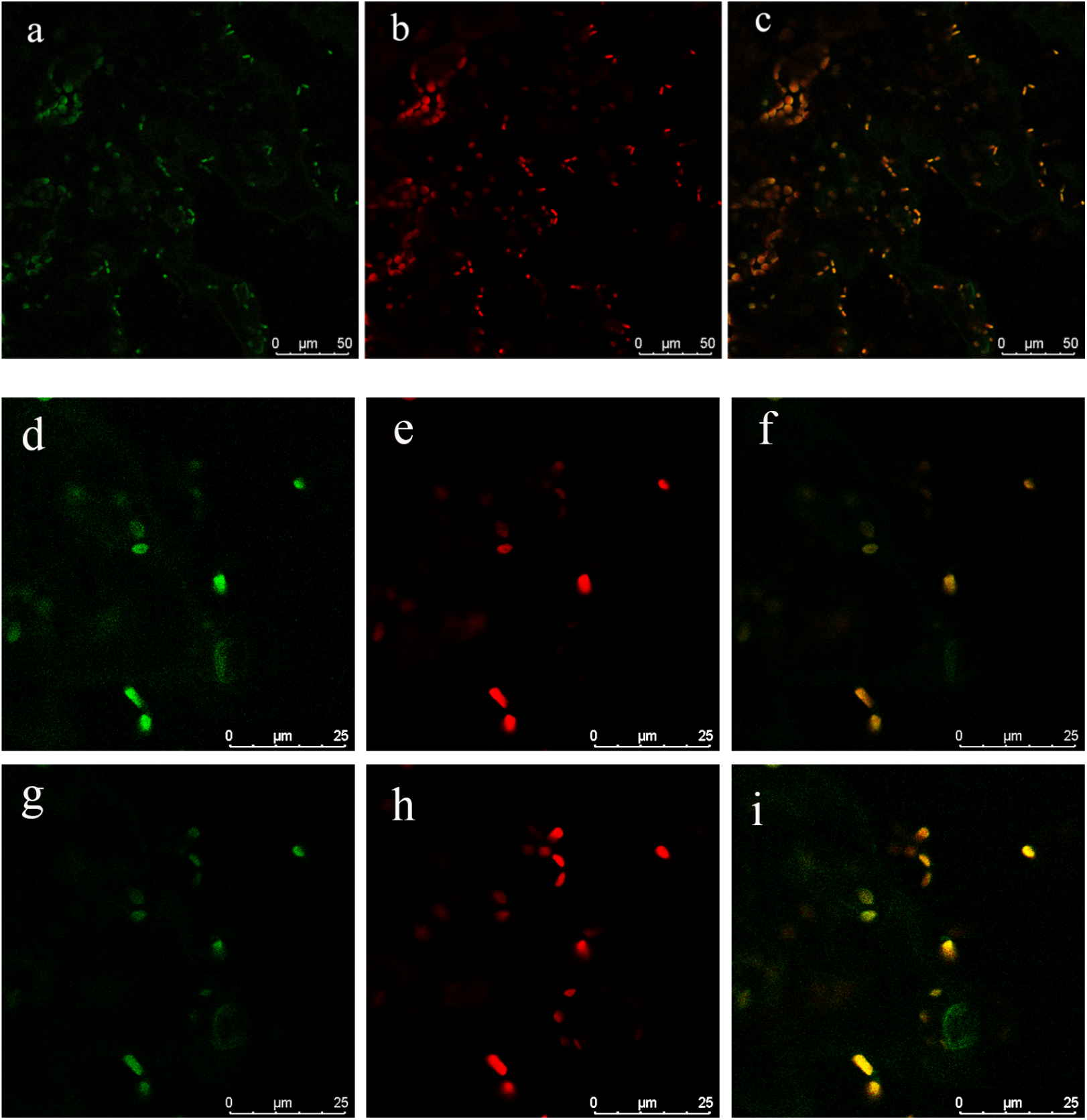
Confocal analysis of stably expressed AIH.1-GFP in *A. thaliana* leaves. Figure (a, b, c- Z-stack), Figure (d, e, f, g, h, i- Z-slices). (a, d, g) GFP-fluorescence (b, e, h) Chlorophyll auto-fluorescence false-colored red (c, f, i) Merged Image.

**Figure 8.**
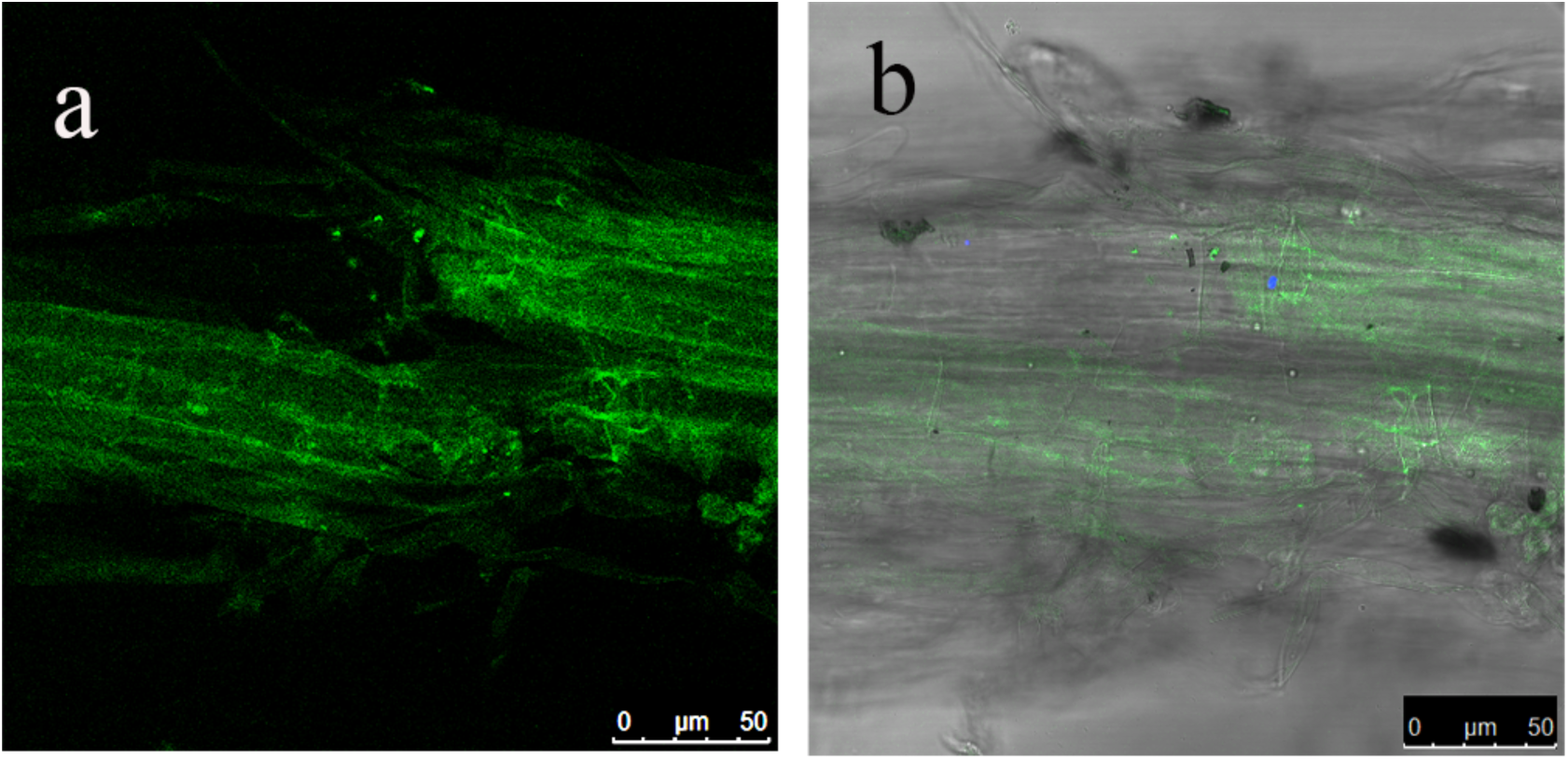
Confocal analysis of stably expressed AIH.1-GFP in *A. thaliana* roots. a) GFP-fluorescence b) Merged image with Brightfield.

### Arabidopsis N-carbamoylputrescine amidohydrolase (AtNLP1) is localized to the ER

N-carbamoylputrescine amidohydrolase1 is the third enzyme in Pathway-2 for putrescine synthesis in plants. NLP1 converts N-carbamoylputrescine to putrescine. There are three isoforms of NLP1 in *A. thaliana*. The longest isoform is NLP1.2 (326aa) long, and the other two isoforms are NLP1.1 (299aa) and NLP1.3 (220aa) (**Figure S.1.2**.). The N-terminal start site for isoforms 1 and 2 is the same. *NLP1.2* has eight exons in common with *NLP1.1* and shares the same N-terminal start site. The shortest isoform *NLP1.3* is missing three exons that are present in the other two isoforms.

*Agrobacterium*-mediated transformation of NLP1.1-GFP and mCherry ER demonstrated that NLP1.1 was localized to the ER. Examination of the Z-stack and individual optical slices showed that GFP expression overlapped the distribution of mCherry in the cell indicating an ER localization of this protein (**Figure 9; Movie S8**). Fluorescence of NLP1.2-GFP also overlapped with the mCherry signals in the cisternal ER (**Figure 10; Movie S9**).

**Figure 9.**
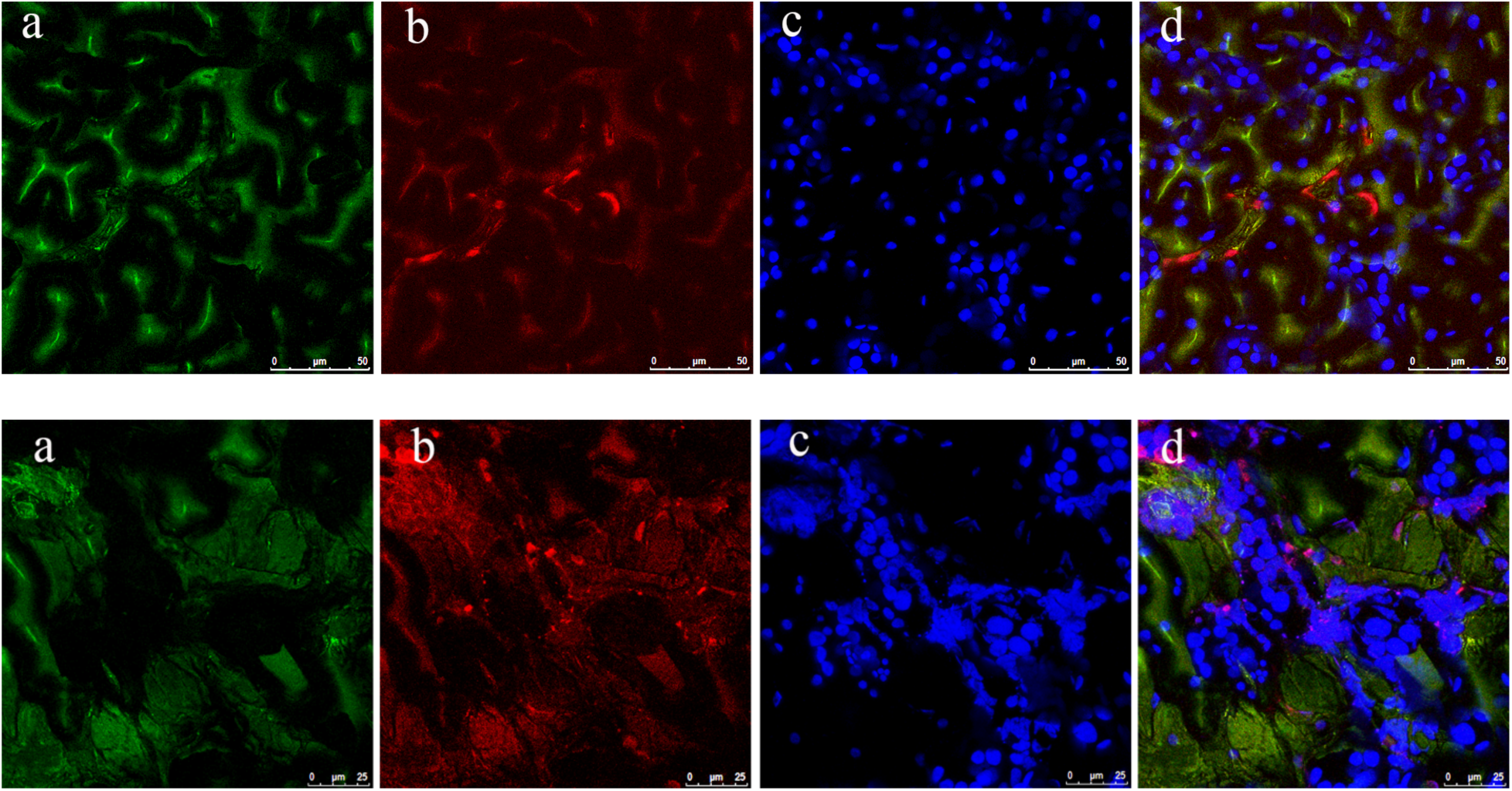
Subcellular localization of AtNLP1.1in the mesophyll cells of *N. benthamiana*. Co-transformation of *AtNLP1.1* fused with GFP along with an ER mcherry organellar marker showing that the gene is localized to the ER. Figure (a, b, c, d - stacked image) a) AtAIH-GFP (b) m-Cherry ER marker (c) Chlorophyll auto fluorescence (d) Merged Scale bars 25μm. Single optical 0.25μm slice of e, NLP1.1-GFP; f, mCherry-ER; g, chlorophyll autofluorescence false colored blue; h, merged image. Fluorescent intensity was uniformly enhanced to facilitate visualization.

**Figure 10.**
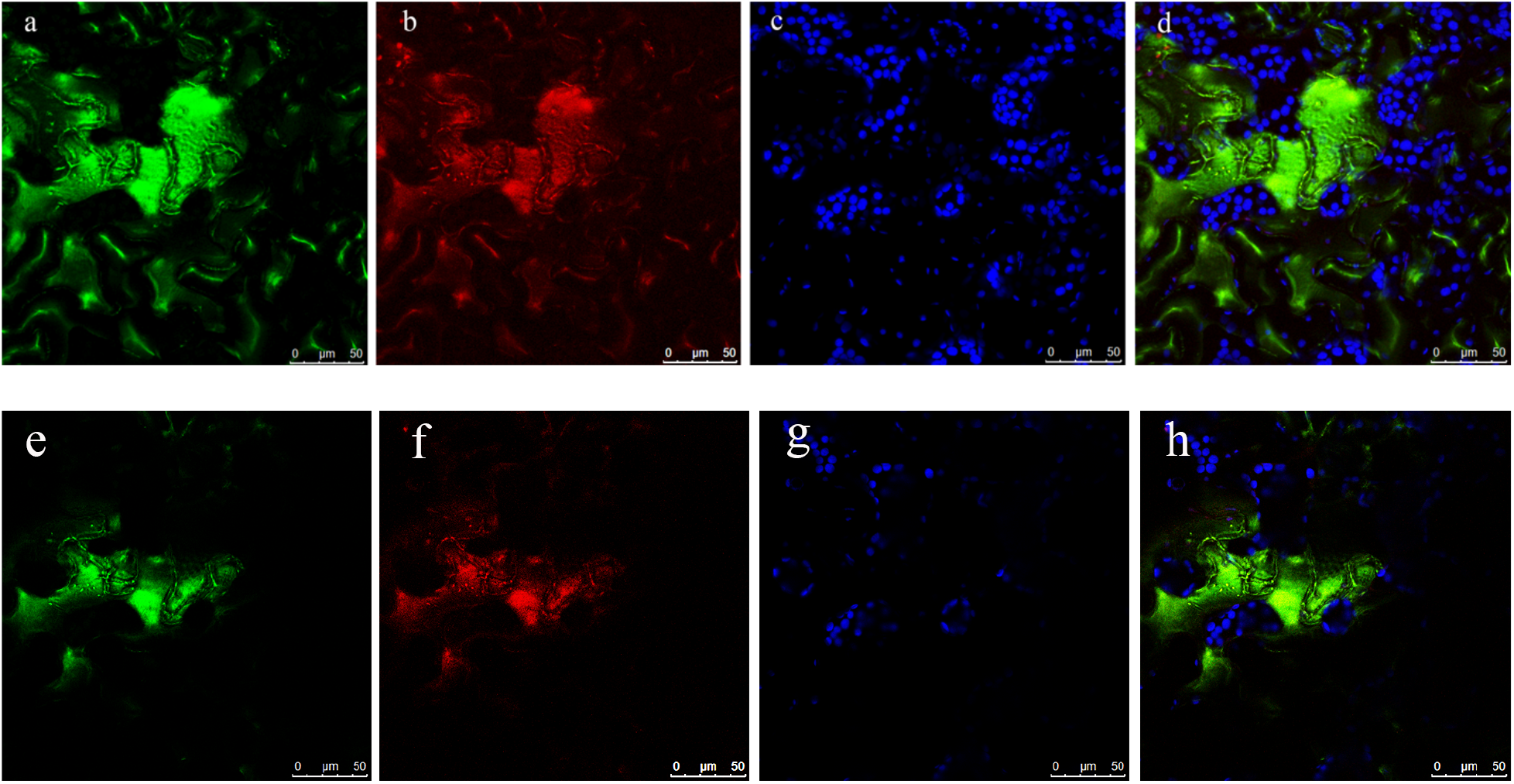
Subcellular localization of AtNLP1.2 in the mesophyll cells of *N. benthamiana*. Co- transformation of AtNLP.2 that was fused with GFP along with an ER mcherry organellar marker showing that the gene is localized to the ER. Chloroplast and ER GFP signals in different areas of the leaf tissue. Figure (a, b, c, d- Z-stack) Figure (e, f, g, h, single optical frame; (a, e) AtAIH-GFP (b, f) ER mCherry marker (c, g) Chlorophyll auto fluorescence (d, h) Merged image. Scale bars 50μm.

BLASTP analysis of AtNLP1.1 against other plant NLPS shows that sequence conservation occurs along the full length of the proteins. NLP1.3 is ~77 amino acids shorter than most other NLPs. The absence of this peptide sequence may indicate that this protein cannot metabolize N-carbamoyl putrescine. Finally, we demonstrated that the co-expression of the shortest isoform- NLP1.3 was also localized in the ER (**Figure 11; Movie S10**). Next, we examined the leaves and roots of a stable transgenic line expressing the longest form AtNLP1.2-GFP (**Figure 12 & 13; Movie S11&S 12**). Confocal analysis of Z-stack images of leaf mesophyll cells showed that GFP fluorescence overlapped with that of chloroplasts along with discrete regions of fluorescence that did not overlap with the chloroplasts (**Figure 12; a, b, c**). To further investigate this association between GFP and chlorophyll fluorescence, we looked at many images from a single optical plane (**Figure 12; d, e, f**). Merged image of the 0.5μm optical slice indicated that fluorescence is adjacent to chlorophyll and is consistent with localization to the ER. In roots, GFP was localized along cell membrane edges displayed as a discrete localization with the cytoplasm (**Figure 13)** conserved with that of the ER localization (Stefano et al.,2015). In summary, stable, and transient expression assays of leaf and root tissues confirmed that all NLP1 isoforms are localized to the ER.

**Figure 11.**
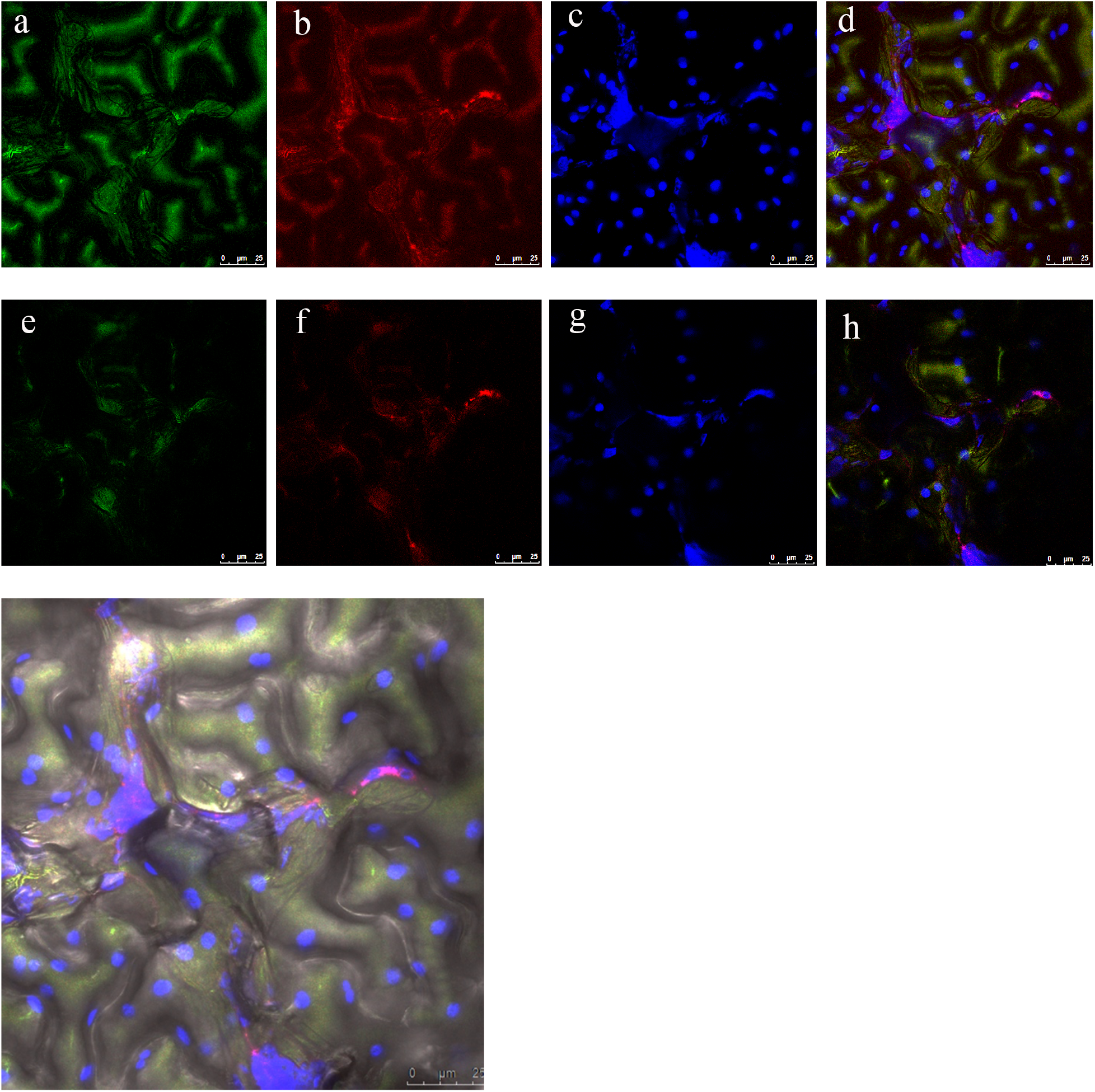
Subcellular localization of NLP1.3 in the mesophyll cells of *N. benthamiana*. Co-transformation of AtNLP1.3 that was fused with GFP along with an ER mCherry organellar marker showing that the gene is localized to the ER. (a, b, c, d: Z- stack images) & (e, f, g, h: Z - slice) (a, e) NLP1.3-GFP (b, f) m-cherry ER marker (c, g) Chlorophyll auto fluorescence (d, h) Merged Scale bars 25μm. Figure (i): Brightfield Image showing GFP signal in the ER.

**Figure 12.**
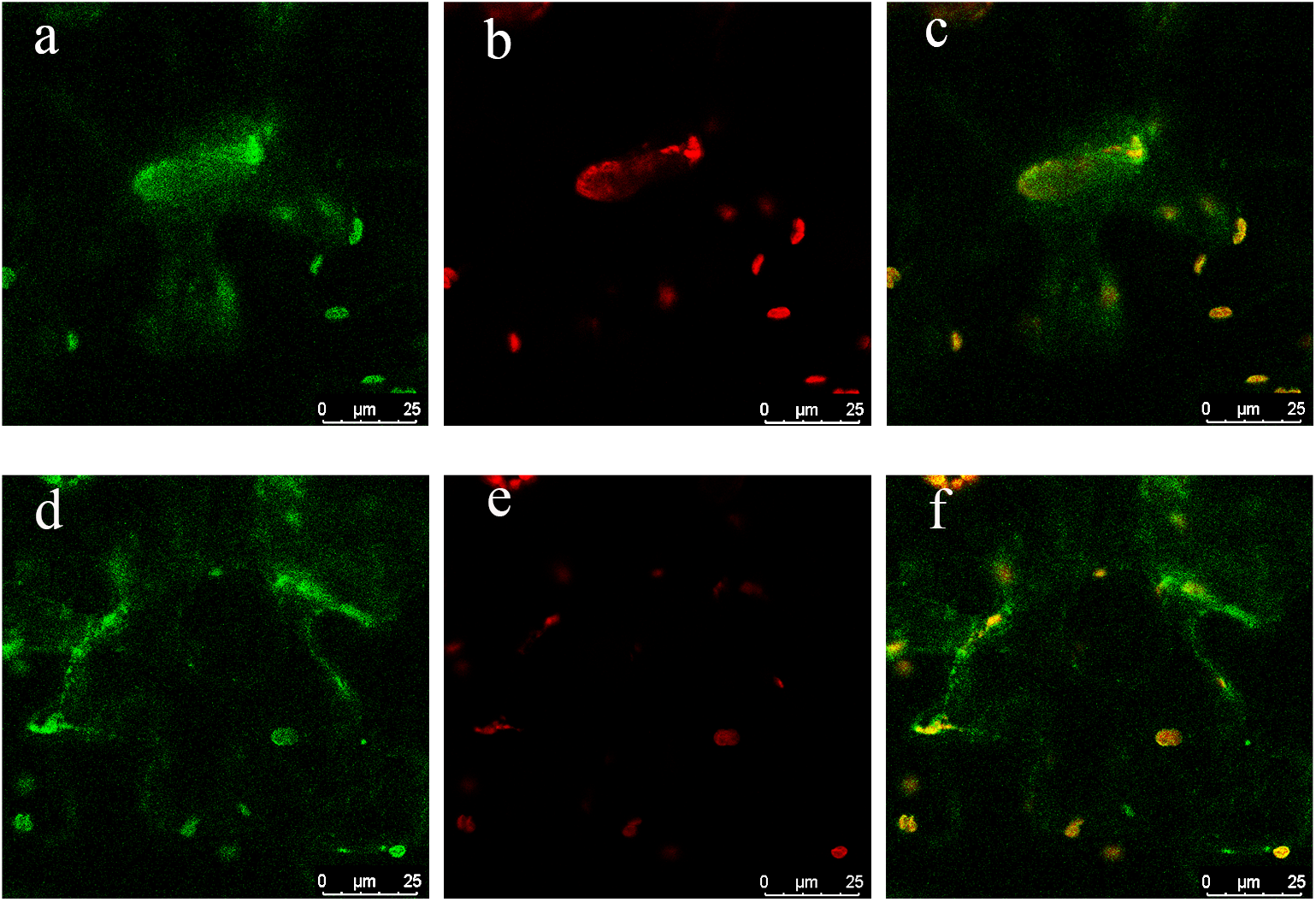
Confocal analysis of stable transformants of NLP1.2-GFP in *A. thaliana*. The leaves from stable transformants were visualized under confocal microscope to determine the subcellular localization of *NLP1.2* gene in *A. thaliana*. Figure (a, b, c Z-stack), Figure (d, e, f- Z-slices) (a, d) GFP-fluorescence (b, c) Chlorophyll auto-fluorescence (c, f) Merged Image shows an ER localization of NLP1.2 gene in the leaves of transgenic *A. thaliana*.

**Figure 13.**
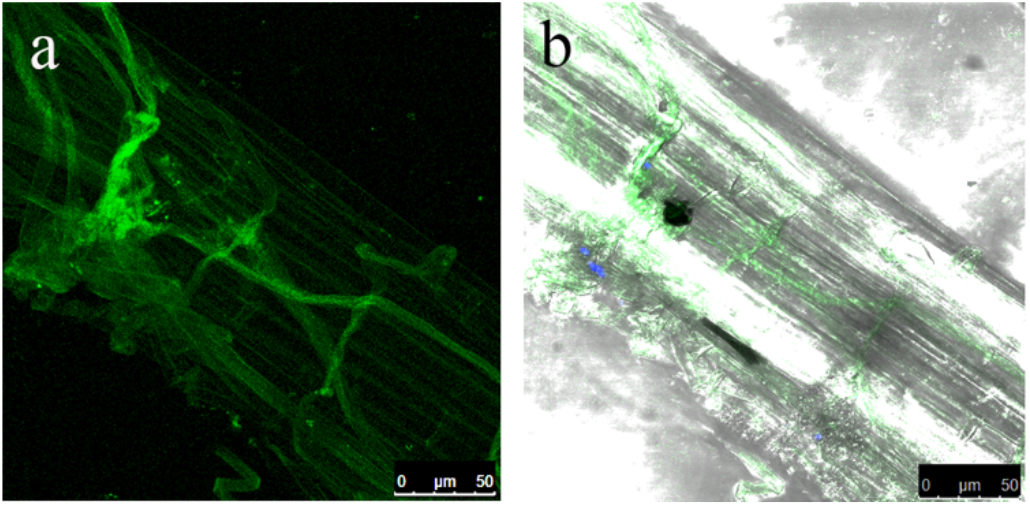
Confocal analysis of stable transformants of NLP1.2-GFP in *A. thaliana*. Representative Z-stacked images of root cells shows that the fluorescent protein is localized to membrane-like structures. a) GFP-fluorescence b) merged image with brightfield

### AIH and NLP genes do not contain conserved uORFs

The 5’UTR regions of plant cells may contain uORFs that function to regulate translational expression (von Arnim et al., 2013). Analysis of uORFs that we extracted from TAIR for agmatine iminohydrolase (AIH) showed that all the three isoforms AIH.1, AIH.2 and AIH.4 contain potential uORF in the 5’ UTR region of the gene. For *NLP1* gene, we found that NLP1.3 contained uORFs in the 5’ UTR region (**Supplemental Table 2,3**). However, BLAST analysis of these uORFs didn’t show any sequence conservation across species. Thus, we concluded that uORFs for AIH.1, AIH.2, AIH.4 and NLP1.3 are unlikely to have a regulatory function

## Discussion

To address the challenges associated with population growth and climate change, plant breeding efforts must identify new strategies to increase crop growth and productivity. Molecular tools that include CRISPR now enable plant biologists to re-engineer plant metabolism (Sweetlove et al., 2017). However, we need to know where metabolites are synthesized within the cells (Lunn et al., 2007). Putrescine is a key signaling molecule that regulates plant responses to abiotic and biotic stresses, and thus a useful target for manipulation by plant breeders. Prior research in our lab showed that manipulation of altered expression of polyamine transporters had profound effects on plant growth and development (Ahmed et al., 2016), but these experiments did not address where putrescine was synthesized and stored. Here, we have used stable and transient expression to localize the key enzymes involved in putrescine synthesis in plant cells and created a model describing the compartmentation of putrescine synthesis in photosynthetic tissues **(Figure 14)**.

**Figure 14.**
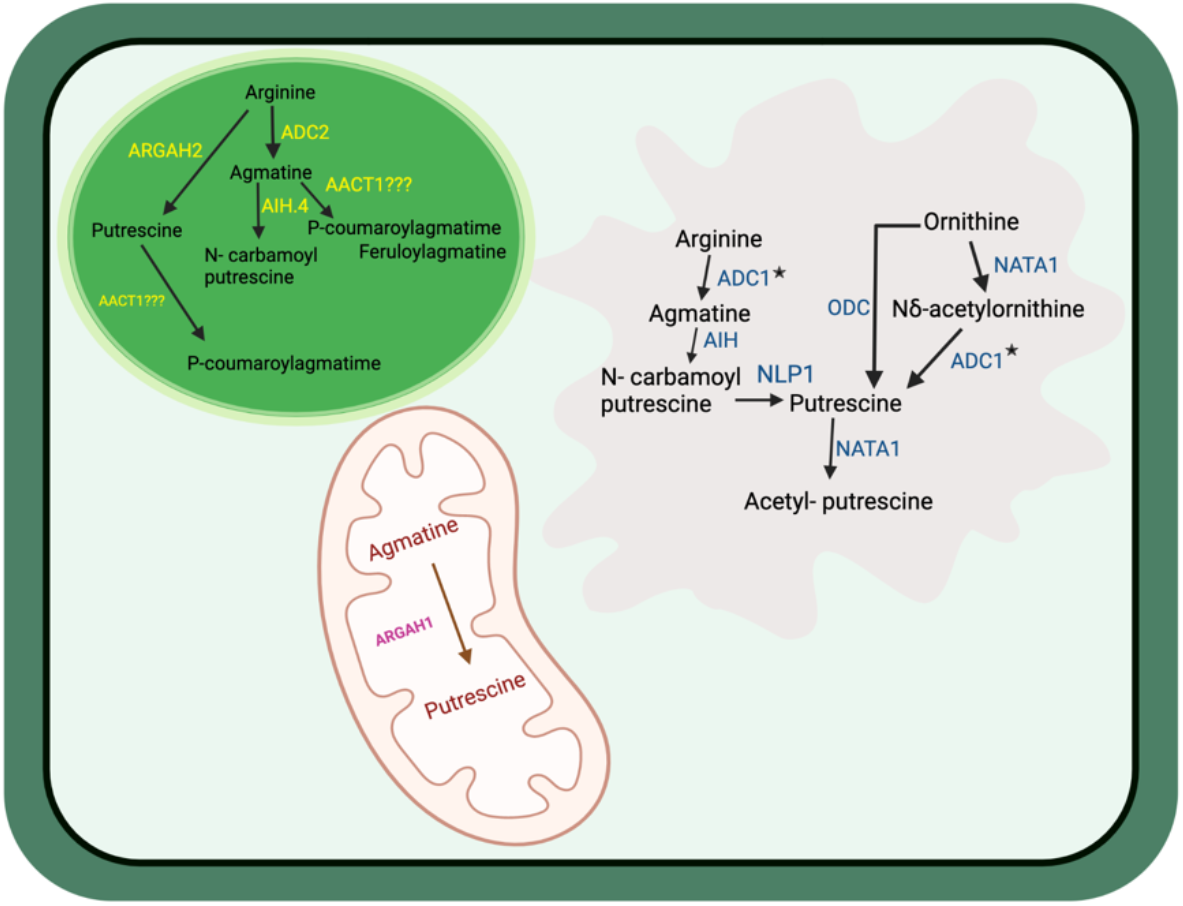
Model illustrating the localization of biochemical pathways for putrescine synthesis and its conjugates in plant cells. Organelles involved in putrescine synthesis include the chloroplast, ER, and mitochondria

The soybean and rice genomes each have two highly similar *ODC* gene models with conservation of sequence at both the N and C terminal regions. Thus, it is expected that the other ODC in these two genomes will also be localized to the ER. ODC’s belong to a highly divergent family as the two proteins; GmODC (Glyma_04g020200.1) and OsODC (LOC_Os04g01590) are only 56% identical and share 70% similarity. Since we observed localization to the ER in both a monocot and a dicot species, we surmise that the ER is a likely site for putrescine synthesis via ODC.

Plant agmatinases (ARGAHs) are strongly conserved across species (Patel et al., 2017; Sekula et al 2020). Previous work from our lab has shown that plant agmatinases have both arginase and agmatinase activity but have a much higher affinity for agmatine so substrate availability of arginine and agmatine, results in the enzymes acting principally as agmatinases (Patel et al., 2017). Here, we observed that rice agmatinase OsAg (*LOC_Os04g01590*) is localized to the mitochondria in leaves. In photosynthetic tissues ADCs have been localized to the ER (Lou et al., 2020) and the chloroplast (Bortolotti et al 2004; Borell et al. 1995; Patel et al 2017). If agmatine that is synthesized elsewhere in the cell, can be imported into the mitochondria, this organelle may also be a significant site for putrescine synthesis.

To address the localization of AIH and NLP1, we made stable C-terminal GFP transformants of the longest isoforms of these proteins in *A. thaliana and* examined the localization of each of the unique protein isoforms of AIH and NLP1 in *N. benthamiana*. Transgenic plants expressing full-length genes for *AtAIH* and *AtNLP1* showed ER localization in both leaf and root tissues. Four of the isoforms of AIH and all the NLP isoforms are also localized to the ER.

Moreover, we observed that AtAIH.4 was localized to both the chloroplast and ER (**Figure 6**). The mechanism behind how proteins co-exist between two different compartments isn’t fully understood. RB60 an atypical protein disulfide isomerase that contains an N-terminal chloroplast signal sequence along with a C- terminal ER retention signal was the first reported protein showing ER and chloroplast localization (Levitan et al.,2005). Many other plant proteins exhibit dual localization (Carrie et al., 2008) with the first report dating back to 1995 (Criessen et al., 1995). Protein isoforms that are location-specific may have specialized functions or may have a new function (Carrie et al., 2008). The product of AIH in the putrescine synthesis pathway is N-carbamoylputrescine, a more reactive substrate than putrescine itself. Thus, in *A. thaliana*, the presence of ADC2 and AIH.4 could result in the synthesis of novel polyamine conjugates that are derived from N-carbamoylputrescine. Synthesis of such conjugates may have critical biological functions. As previously noted, plants with *aih* mutations have an embryo-defective phenotype (Tzafrir et al., 2004). The molecular basis of this phenotype was originally attributed to the loss of putrescine synthesis, but mutations in NLP1 do not have a phenotype because *A. thaliana* has other pathways for putrescine synthesis (Patel et al., 2017; Lou et al., 2020). Thus, the molecular basis of this phenotype is still unknown.

To further demonstrate that AIH.4 contains a plastid signal as predicted by (Kuo-Chen et al., 2010) and confocal imaging, the N terminal sequence can be fused directly to GFP and transformed into *N. benthamiana*. Perhaps the most direct way to test the importance of localization of this isoform to the chloroplast would be to determine if increased expression of AIH.4 produced a novel phenotype.

Differences in the 5’UTR’s and in the coding sequences of the N-terminal region of AIH and NLP1 could result in differential expression in different cell types or differential rates of translation (Reviewed in Chaudhary et al.,2019). Additional regulation could come from uORFs in the 5’ UTRs (Calvo et al., 2009; Lin et al., 2019; Zhang et al., 2020). *AIH.1, AIH.2, AIH.4*, and *NLP1.3* have uORFs but we didn’t observe conservation of the uORFs in the Brassicas. Thus, translational regulation of these genes by uORFs is not known and needs further investigation.

Our findings that the ER is a major site of putrescine synthesis in plants has significant implications for plant metabolism. The localization of the enzymes NATA1 and ADC1 (Lou et al., 2020) and the isoforms of AIH and NLP1 to the ER, may indicate that whenever NATA1 is present, much of the putrescine in the ER is converted to acetyl-putrescine. In plant cells, the ER is the site of synthesis for one-third of the cellular proteome, including membrane receptors, transporters, and cell wall components. Proper transmembrane protein processing is crucial because transmembrane proteins are particularly responsible for plant response to different stresses. Any glitch in this process directly affects plant development and immunity (Eichmann and Schafer 2012). Polyamines have very specific interactions with some membrane transporters (Schuster and Bernhardt 2011) and have recently been shown to mediate the folding of very acidic proteins (Despotovic et al., 2020). Elevated levels of positively charged polyamines may aid in the proper folding and stability of these proteins during vesicular transport. When plants are under stress, cellular demands for the folding of secretory proteins are elevated which sometimes may exceed the ER protein folding abilities. This results in an accumulation of unfolded proteins in the ER, and activation of the unfolded protein response pathway (UPR) (Eichmann and Schafer 2012). Could decreased putrescine availability in the ER trigger an enhanced UPR?

The plant ER transverses across cells via plasmodesmata creating a tissue-spanning ER network. A single ER network throughout the plants connecting cells via desmotubules may regulate metabolite flow between cells (Henne 2021; Brandizzi 2021). Thus, putrescine that is synthesized in the ER can be exported from cell to cell via the plasmodesmata. Putrescine is the direct precursor for higher polyamines thermospermine, spermidine, and spermine. Synthesis of higher polyamines requires S-adenosyl methionine and results in the production of 5-methylthioadenosine. The Yang cycle includes the enzymes that recycle 5-methylthioadenosine to methionine in plants, and these enzymes appear to be restricted to the vascular tissues (Pommerrenig et al., 2011). Consistent with the localization of the Yang cycle, the synthesis of thermospermine is primarily localized to phloem tissues (Clay et al., 2005) and in potatoes, spermidine synthase is also restricted to phloem tissues (Sichhart and Drager 2005). The ER localization of two putrescine synthesis pathways in plants may facilitate the regulated transfer of putrescine to phloem tissues via plasmodesmata.

The localization of putrescine synthesis to the ER could also be a strategy to facilitate either the export of putrescine from the cell or its trafficking to the vacuole without altering cytoplasmic concentrations of putrescine. Perhaps, the most direct way to assess the role of the ER in putrescine export would be to generate mutants in ADC1 and NLP1 and to determine whether mutations in ADC1 reduced polyamine in the apoplast via the phloem to other tissues. The secretion pathway originating in the ER and going from the Golgi to the plasma membrane may be a mechanism for the delivery of both proteins and small molecules such as polyamines to the apoplast (Poustka et al., 2007).

The ER is a dynamic network of tubules and cisternae with contact sites between the plasma membrane, nucleus, Golgi, and the chloroplast (Friedman and Voeltz 2011). In the transgenic localization data for AtAIH.1, GFP signal appears in the ER regions of the roots and leaves. There are areas on the chloroplast where the GFP signal overlaps with a portion of the chloroplast in what appears to be ER chloroplast contact sites. The pattern of GFP localization in the ER and the chloroplast contact sites is similar to images captured by (Andersson et al., 2007). ER- chloroplast membrane association may facilitate the transport of lipids and other cellular metabolites that have precursors in the chloroplast (Andersson et al., 2007). In these experiments, *Arabidopsis* protoplasts were first ruptured using laser scalpel to release cell contents. In the cell lysates, ER fragments remained attached to the chloroplast and couldn’t be separated from one another, suggesting a strong protein-protein interaction. (Tan et al., 2011) showed that CLIP1 lipase/acylhydrolase (BnCLIP1) in *Brassica napus* is localized at the ER-chloroplast contact sites in *N. benthamiana*. Polyamines are charged molecules, so exchange between the ER and the chloroplast must take place via membrane transporters, but close apposition of these membranes could facilitate the transfer of agmatine synthesized by ADC2 in the plastid for synthesis to putrescine via AIH and NLP1. Regions within the cell where the chloroplast and ER membranes are very closely aligned may facilitate the exchange of nonpolar metabolites without the need for specific transporters. Polyamines synthesized in the ER could potentially also be transferred directly to the vacuoles, which has already been demonstrated for anthocyanins transport (Poustka et al., 2007).

As previously noted, little is known about how plant cells sense and respond to changes in polyamine levels (Kusano et al., 2018). However, one of the observations that we have made is that changes in the expression levels of polyamine transporter in the PUT family have pleiotropic effects (Ahmed et al., 2017). Members of the PUT family of polyamine transporters function as unidirectional putrescine and spermidine transporters. The *A. thaliana* genes *PUT2* and *PUT3*, and the rice gene *OsPUT3* have been localized to the chloroplast, while both *AtPUT5* and the rice gene *OsPUT1* have been localized to the ER (Ahmed et al.,2017). Overexpression of *PUT3/RMV1* enhances the toxicity of paraquat (Fujita et al., 2012), and thus is assumed to be oriented to pump polyamines into the chloroplast. The role of PUT transporters such as OsPUT1 or AtPUT5 which are localized to the ER transporters is not fully understood. However, the overexpression of either chloroplast localized PUTs, which pump polyamines into the chloroplast from the cytosol; or ER-localized PUT transporters have similar effects on leaf size, time to flowering, and floral stem thickness (Ahmed et al., 2017). These results suggest that altering the availability of polyamine levels in the cytoplasm results in changes in gene expression affecting developmental pathways.

Our findings indicate that putrescine synthesis in plants is specifically excluded from the cytoplasm and is tightly regulated by polyamine transport. This observation contrasts with that of humans where ODC is localized to the cytoplasm and the nucleus (Schipper et al., 2004).

The exclusion of putrescine synthesis from the cytosol and the differential activation of different pathways may allow putrescine to function as a signaling molecule that alters the expression of hormone regulated pathways. Increasing putrescine levels mediate cold and freezing tolerance in part by increasing ABA synthesis (Cuevas et al; 2008). Both ADC1 and ADC2 contribute to freezing tolerance, but ADC1 is activated much earlier than ADC2 (Cuevas et al; 2008). This could indicate that the location where putrescine is synthesized, is also important in integrating the plant response to stress. Overexpression of ADC2 resulted in dwarfism and late flowering due to inhibition of GA synthesis (Alcazar et al., 2005). Increased expression of PUT transporters in *A. thaliana* resulted in a delay of flowering, but a two-fold or more increase in floral and seed production (Ahmed et al., 2017). Putrescine levels were shown to affect root meristem size by modulating auxin signaling (Hashem et al.,2021). Catabolism of putrescine is also linked to the synthesis of GABA another plant signaling molecule via copper amine oxidases and NO (Wuddineh et al., 2018; Zarei et al., 2016).

Our study provides the understanding of the different pathways mediating putrescine synthesis and some of its conjugates in photosynthetic tissues (**Figure 14)**. The enzyme AACT1 was included because it is significantly upregulated in biotic and abiotic stresses and its preferred substrates are agmatine and putrescine (Winter et al., 2015). An obvious deficit in our present understanding of putrescine metabolism is how substrates are mobilized within the cell itself. We hypothesize that the myriad of phenotypic changes attributed to putrescine can be traced back to the activation of specific pathways. The extent of crosstalk between pathways and the relative importance of pathways in response to biotic and abiotic challenges will need to be addressed. This will require genetic analysis of knockouts of key genes in each of the pathways combined with transcriptomics, proteomics, and metabolomics. The potential role of ER in the synthesis and export of putrescine and putrescine conjugates also needs to be addressed.

